# CAMSAP3 loss of function models suggest causative role in generalized genetic epilepsy

**DOI:** 10.64898/2026.09.01.744686

**Authors:** Christopher Mark LaCoursiere, Zachary Stayn, Hannah Hepner, Sneham Tiwari, Joseph Pascucci, Chariton Moschopoulos, Lacey Smith, Hyun Yong Koh, Parul Chaudhary, Annapurna Poduri

## Abstract

Advancements in next-generation sequencing have led to the discovery of hundreds of human epilepsy gene associations. Newly associated genes require functional validation to establish causation and to inform patient treatment in the clinic. A recent exome sequencing trio analysis identified predicted protein-altering variants in two patients with generalized epilepsy in the gene *CAMSAP3*. *CAMSAP3* regulates non-centrosomal microtubule dynamics and the acetylation necessary for normal axonal differentiation and migration. We show that overexpression of patient variants leads to protein degradation and dysregulation of microtubule acetylation in cultured HEK cells. *Camsap3* knockout zebrafish also exhibit increased axonal microtubule acetylation, as well as epileptic features such as seizure-like swimming behaviors, aberrant inhibitory interneuron development, and epileptiform via electrophysiology. Together, these data suggest that *CAMSAP3* plays an important role in genetic generalized epilepsy.

**Highlights:**

- Loss-of-function *CAMSAP3* variants in individuals with genetic generalized epilepsy alter tubulin dynamics in HEK cells.
- *Camsap3* loss-of-function zebrafish show dysregulation of acetylated tubulin in tectal axon projections.
- Mutant fish exhibit hyperexcitable swimming behavior, loss of inhibitory interneurons and epileptiform via electrophysiology

## Introduction

Epilepsy is a neurological disorder defined by recurrent unprovoked seizures due to increased, abnormal electrical activity in the central nervous system (CNS). This disorder impacts roughly 1% of the global population.^1^ Despite this high prevalence, many individuals’ epilepsies are classified as idiopathic.^2^ Seizures are generally classified into two broad categories, focal and generalized, based on their onset location as identified by electroencephalogram (EEG). Focal seizures are localized to a single area of the brain, while generalized seizures occur simultaneously on both sides of the brain. Generalized seizures include absence seizures, which feature episodic loss of consciousness; myoclonic seizures, characterized by involuntary muscle jerks, typically in the arms, legs and torso; and tonic-clonic seizures, which feature stiffening of the body followed by rhythmic jerking.^3^

Generalized epilepsy disorders have numerous causes, including genetic abnormalities.^2^ Genetic generalized epilepsies (GGE) often present early in childhood and can carry significant risks, including status epilepticus, in which prolonged seizure activity can lead to neuronal injury and other long-term consequences; sudden unexpected death in epilepsy (SUDEP), in which a person with epilepsy dies without a known cause of death; and changes in memory and/or cognition due to seizure-induced interruptions of normal brain function.^4–6^ Identifying a genetic etiology has been shown to mitigate these risks and improve long-term outcomes, in part due to changes in management based on the specific genetic diagnoses identified.^7,8^ Further, early diagnosis allows for early interventions, including physical and occupational therapy, speech and feeding therapy, and learning support. One recent cohort study revealed two *de novo* variants, one frameshift with premature truncation and one missense, in the *CAMSAP3* gene in two patients with genetic generalized epilepsy (GGE).^9^

CAMSAP3 is essential for anchoring and dynamically inhibiting tubulin acetylation in non-centrosomal microtubules.^10^ Microtubule abnormalities are a common contributor to neurodevelopmental disease^11^ because of their crucial role in neuronal developmental processes such as cell migration and axonal differentiation.^12^ Differentiation of nascent axons are determined by the accumulation of stabilized and mature microtubules within a single neurite. This is because the development of neuronal polarity depends on a positive feedback loop, driven by cargo shuttling along stable microtubules. Within a nascent axon, dendrite differentiation signals are shuttled to distal neurites and axon differentiation signals are shuttled throughout the nascent axon along stabilized microtubules. Tubulin acetylation is a post-translational modification necessary for stabilizing microtubules and regulating this feedback loop. *Camsap3* knockdown in cultured mouse hippocampal neurons leads to an increase in microtubule acetylation and thus, an increase in stabilized microtubules within multiple developing neurites. Multiple neurites containing mature, stabilized microtubules leads to a loss of polarity, which leads to the development of supernumerary axons. While there is robust evidence to support CAMSAP3’s role in neuronal development, its relevance *in vivo* and its connection to human neurological disease have not been fully elucidated.

To determine whether the *CAMSAP3* variants identified in human patients contribute to their epilepsy, we designed a zebrafish model expressing a truncating *camsap3* variant similar to that which was previously reported ^9^ and evaluated for epilepsy-related phenotypes. We evaluated this *camsap3* loss-of-function (LOF) model for the following features: 1) seizure-like behavior, as measured by swimming parameters such as convulsions, whirlpool behavior, and posture loss events; 2) abnormal electrical bursting and spiking in certain regions of the brain, as assessed through local field potential (electrophysiology) recordings; 3) dlx-positive inhibitory interneuron number and organization; and 4) abnormal fish brain and body morphology. In addition, we evaluated HEK cells overexpressing each of the two reported patient CAMSAP3 variants for altered tubulin dynamics. Here, we present evidence that *camsap3* loss of function causes behavioral seizures, abnormal electrical activity, and loss of inhibitory interneurons in zebrafish, as well as cellular evidence that specific variants alter tubulin acetylation dynamics. Together, these lines of evidence support a role for *CAMSAP3* in neurodevelopment and establish it in the pathogenesis of epilepsy.

## RESULTS

### *CAMSAP3* variants in two patients with generalized epileptiform EEG patterns

We first sought to delineate the phenotypes associated with *CAMSAP3* variants in two children with epilepsy reported in Koh, Smith et al ^9^ (Figure 1). The two patient variants are located in sub-domains of the CAMSAP3 protein that are both involved in microtubule binding. Patient 1 has a heterozygous frameshift variant (c.3507del; p.(K1170RfsX17)) located in the CKK region, resulting in premature termination. Patient 2 has a heterozygous missense variant (c.3068A>C; p.(K1023T)) located in the C-terminal D2 domain. *In silico* annotation for the CAMSAP3 p.(K1023T) variant yielded the following predictions: PolyPhen = 0.996 (probably damaging), CADD = 23.4 (highly pathogenic), and SIFT = 0.21 (tolerated). While these predictor outputs were mixed, the high CADD and PolyPhen metrics strongly suggest that the variant is deleterious. Gene-wide constraint metrics from population data within gnomAD demonstrated a high probability of loss-of-function intolerance (pLI=1). Missense constraint scores were mixed, with a gene-wide gnomAD missense Z-score of 1.71 while regional constraint surrounding the missense variant (19-7615000-7616000) demonstrated a higher Z-score of 6.52. Missense tolerance ratio (MTR) showed higher tolerance with MTR=1.00.

**Figure 1.**
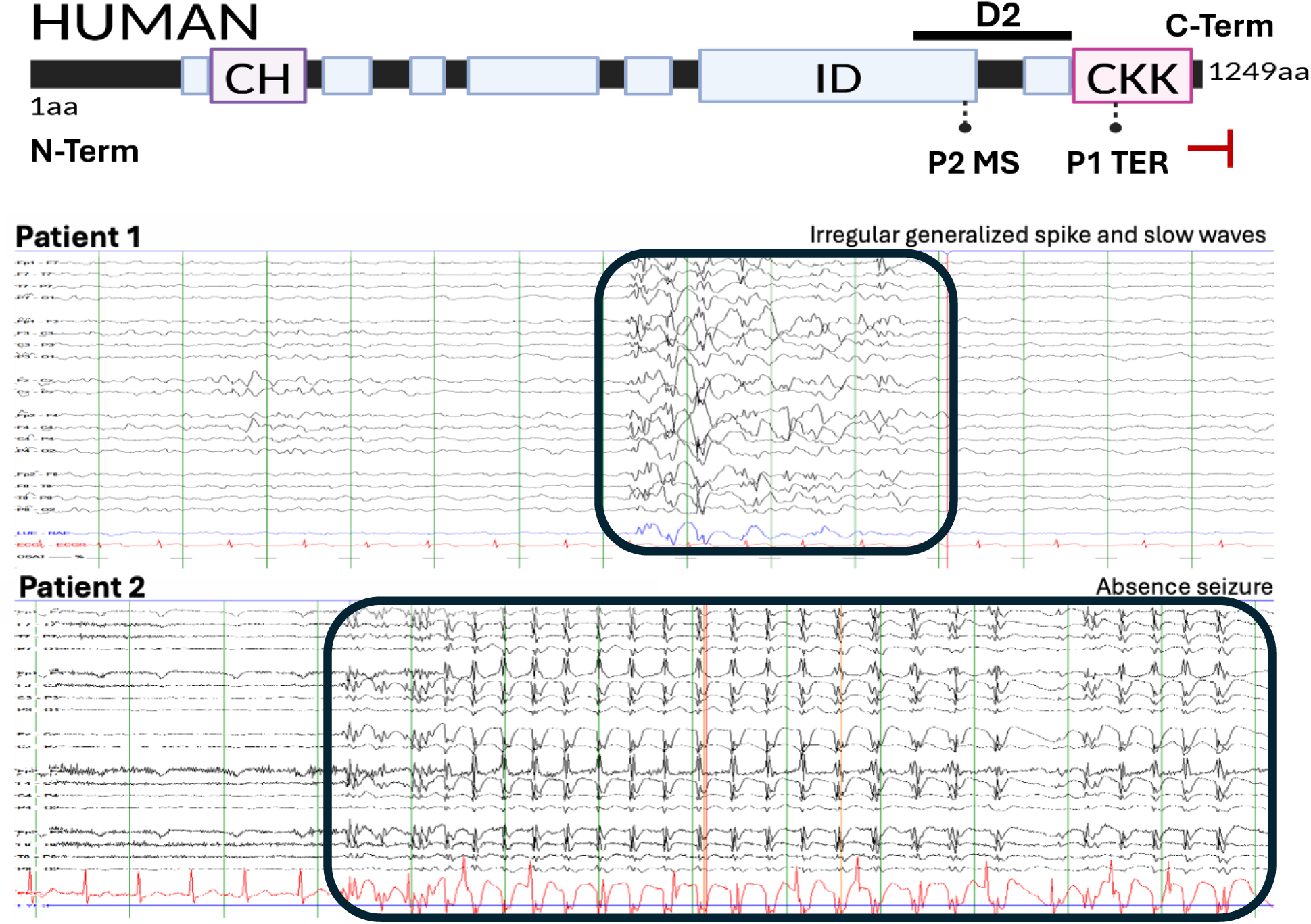
Generalized epileptiform EEG activity in two patients with *CAMSAP3* variants mapping to the D2 and CKK microtubule-binding domains. Each patient’s variant is mapped onto the CAMSAP3 protein. Patient 1’s variant, K1170RfsX17, is in the CKK domain that binds the minus end of non-centrosomal microtubules; the frameshift variant creates a premature truncation (Ter). A sample of his EEG shows rhythmic generalized spike-wave complexes during a facial myoclonic seizure. Patient 2’s missense variant, K1023T, is located in the D2 microtubule expanded lattice binding domain. A sample of her EEG displays rhythmic generalized spike-wave complexes during an absence seizure. (EEG was acquired using the bipolar montage. Vertical green lines represent one-second intervals, and sensitivity of the recordings is 7 µV/mm.)

Patient 1 is a male with seizure onset at 5 years and 2 months who presented with generalized tonic-clonic seizures, absence seizures, and facial myoclonus. EEG showed irregular generalized spike-and-slow-wave discharges during an observed facial myoclonic seizure (Figure 1). He had mild learning disabilities including dyslexia, no family history of epilepsy, and normal MRI. Seizures were controlled with ethosuximide and valproic acid since age 7.

Patient 2 is a female who had seizure onset at 9 years of age; EEG at the time of presentation showed generalized spike-wave complexes during an absence seizure, consistent with a generalized epilepsy diagnosis (Figure 1). She had recurrent absence seizures that progressed to generalized tonic-clonic seizures by age 17 and normal development with no reported learning disabilities or dysmorphic features; her seizures have been controlled with lamotrigine since age 22.

### Patient *CAMSAP3* variants modeled in HEK cells disrupt acetylation of tubulin

To evaluate whether human variants lead to *CAMSAP3* LOF, we overexpressed GFP-tagged *CAMSAP3* (WT) into human HEK-293T cells. Variant containing constructs were generated to contain either the K1023T missense variant (MS) or the frameshift K1170RfsX17 variant (KO) and were validated via Sanger sequencing (Supplemental Figure 1). Though non-neuronal, HEK cells were used because of their high transfection efficiency and because they have been previously used to evaluate the effects of *CAMSAP3* overexpression on tubulin acetylation.

Analysis of GFP intensity in transfected HEK cells revealed genotype-dependent differences in the expression of CAMSAP3. WT and MS overexpressed cells exhibited strong GFP signal compared to non-transfected vehicle controls (VEH) and KO overexpressed cells (Figure 2A). These data demonstrate successful expression of the WT and MS CAMSAP3-GFP constructs, while the frameshift construct showed markedly reduced GFP signal, consistent with premature truncation.

**Figure 2.**
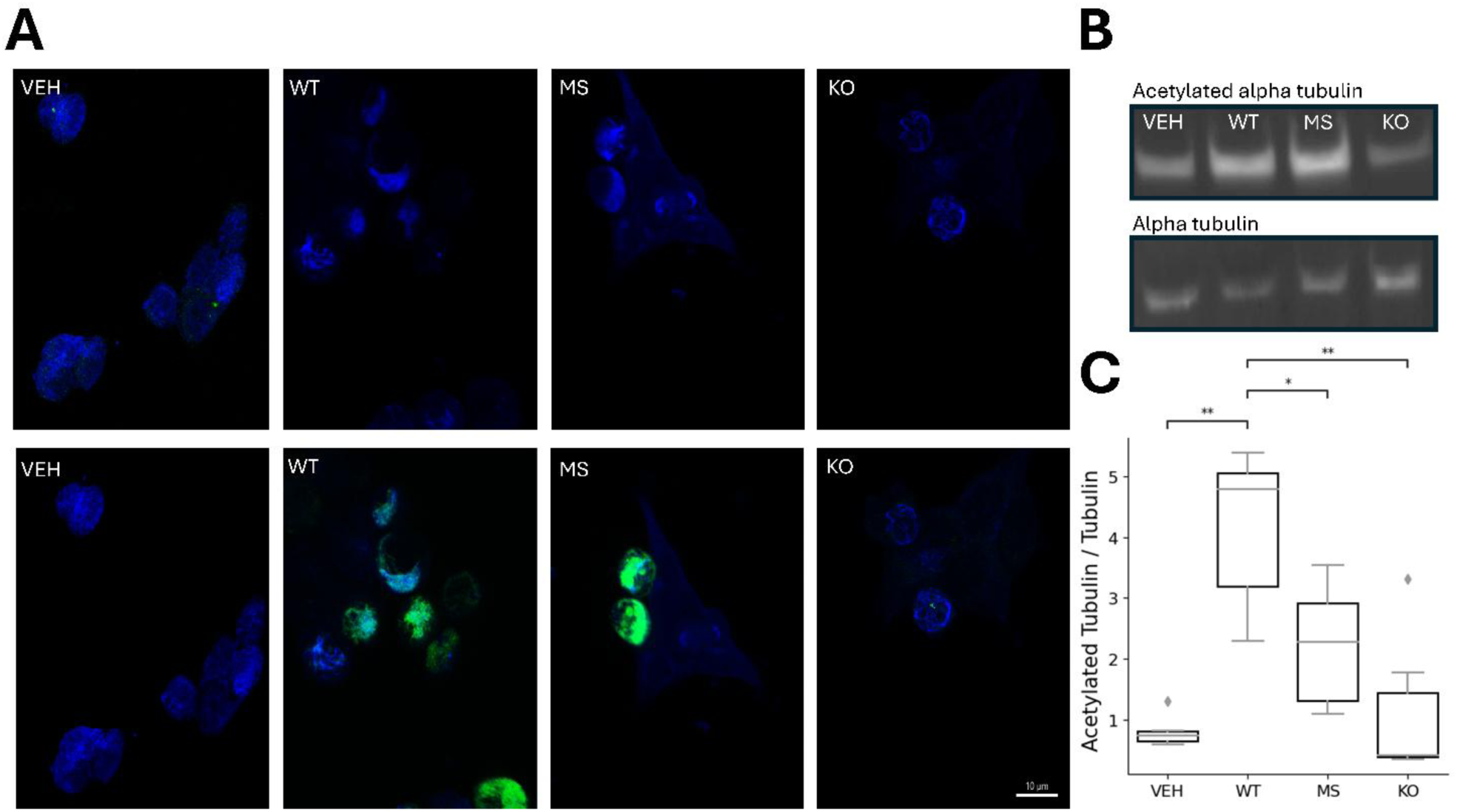
*CAMSAP3* variant overexpression in HEK293 cells disrupts tubulin dynamics. (A) GFP signal is abundant in WT-transfected cells and in cells expressing the *CAMSAP3* gene with the K1023T missense variant (MS). GFP signal is markedly absent in cells expressing CAMSAP3 the K1170RfsX17 variant (KO), consistent with truncation and possible nonsense-mediated decay. (B-C) Overexpression of the WT *CAMSAP3* shows increased acetylated alpha tubulin relative to total alpha tubulin compared to VEH control via western blot. Overexpression of MS and KO *CAMSAP3* leads to a relative decrease in acetylated tubulin (MS:KO p<1x10^-4^, MS:VEH p=0.993, MS:WT p<1x10^-4^, KO:VEH p=0.10, KO:WT p<1x10^-4^, VEH:WT p<1x10^-4^). Significance was determined by ANOVA with Tukey post hoc tests; ns=not significant, *p<.05, **p<.01, ***p<.001, ****p<.0001.

Immunocytochemistry showed significant differences of acetylated α-tubulin across experimental groups (Figure 2A). Cells expressing WT *CAMSAP3* exhibited significantly greater acetylated α-tubulin relative to total tubulin when compared to VEH, MS, and KO transfected groups. This effect was most pronounced in the KO; nearing VEH levels of acetylation. Compared to WT, acetylation was still decreased in the MS group but remained well above levels seen in VEH (Figures 2B-C). Together, these findings demonstrate that supraphysiological overexpression of WT *CAMSAP3* increases acetylation in HEK’s, whereas *CAMSAP3* patient variants results in lower acetylation in HEK cells compared to WT, suggesting these variants lead to LOF

### CRISPR/Cas9-edited zebrafish have *camsap3* loss of function

We generated an out of frame, 4 base pair deletion, in the *camsap3* zebrafish gene using CRISPR/Cas9. The resulting frameshift codes for premature truncation early in the CKK tubulin-binding domain of Camsap3 (p. Asp1270fsTer17) (Figures 3A-B). This early truncation results in nonsense-mediated decay and complete loss of function of the Camsap3 protein. To confirm that this premature truncation reduced *camsap3* expression, we quantified *camsap3* mRNA levels by RT-PCR in the zebrafish knockout model and compared them with those with the wild-type control zebrafish (Figure 4C).

**Figure 3.**
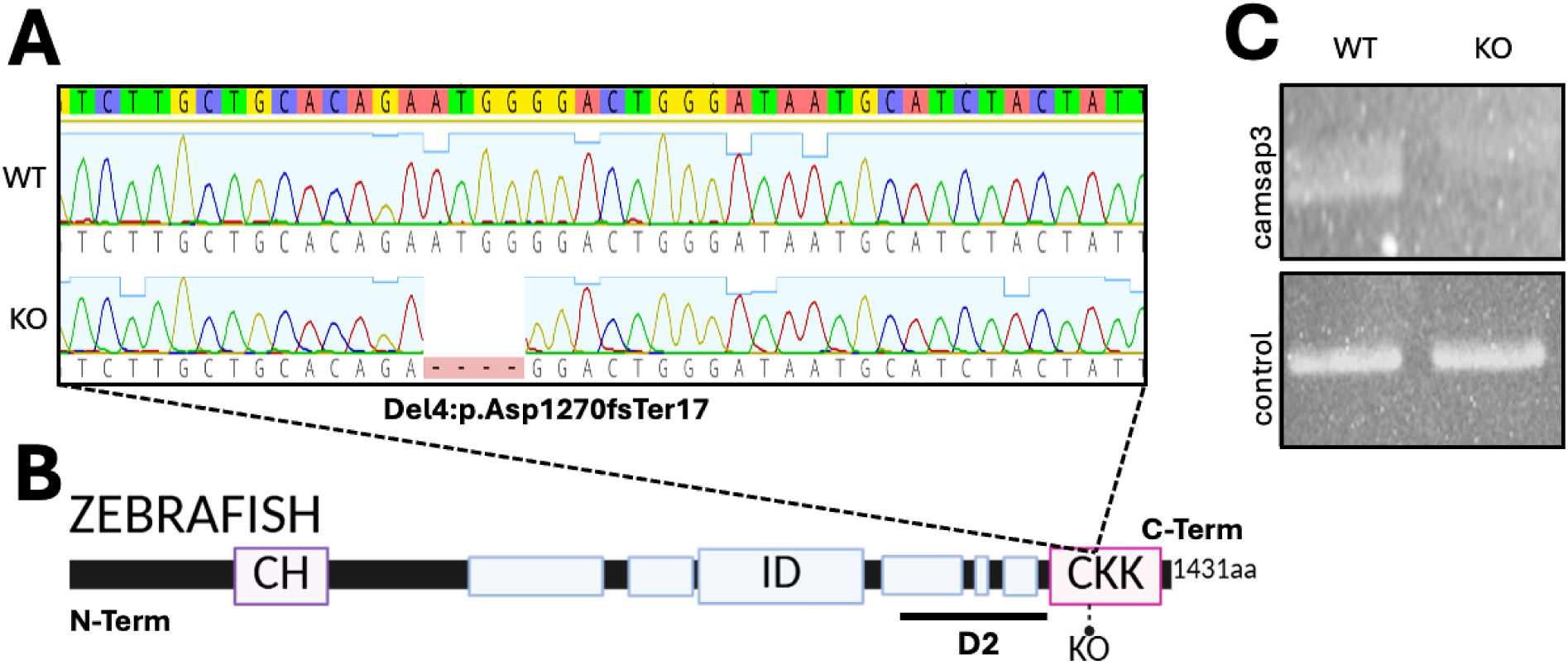
CRISPR/Cas9-mediated gene editing results in a loss-of-function zebrafish model of *camsap3*. (A) Sanger sequencing of 6 dpf larval DNA showing homozygous four-base pair deletion in *camsap3* mutants. (B) Schematic representation of zebrafish Camsap3 protein showing the location of the introduced mutation. This out-of-frame deletion is predicted to result in a frameshift and early truncation within the *camsap3* tubulin-binding domain. (C) N-terminal cDNA amplification of *camsap3* shows that the mutation leads to RNA degradation.

**Figure 4.**
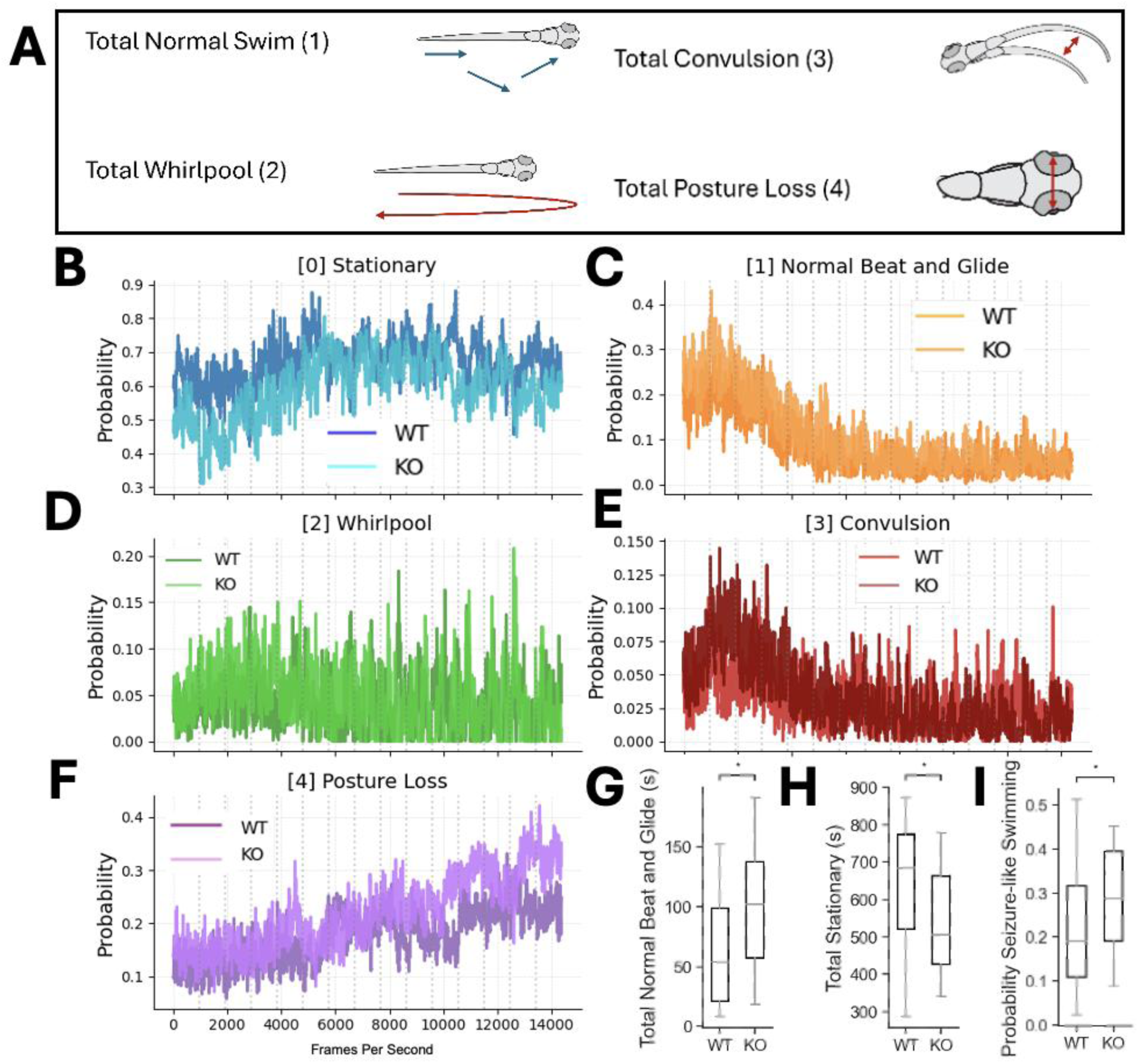
*Camsap3* KO larvae show increased hyperexcitable swimming patterns in response to the proconvulsant PTZ compared to control larvae. (A) Representative swimming behaviors seen in zebrafish wild-type (WT) and knockout (KO) larvae; whirlpool, convulsion, and posture loss behaviors are considered seizure-like behaviors. (B) Difference in stationary behavior between WT (dark blue) and KO (light blue) fish over time; WT and KO larvae have similar median probability of stationary behavior (p > 0.05 at each minute). (C) Difference in beat-and-glide swimming behavior between WT (light orange) and KO (dark orange) larvae over time; WT and KO larvae have similar median probabilities of normal beat-and-glide behavior (p > 0.05 over each minute) (D) Difference in whirlpool behavior between WT (dark green) and KO (light green) larvae over time; WT and KO larvae have similar median probabilities of whirlpool behavior (p > 0.05 at each minute). (E) Difference in convulsion behavior between WT (bright red) and KO (dark red) larvae over time; WT and KO fish have similar median probabilities of convulsion behavior (p > 0.05 at each minute) (F) Difference in total posture loss between WT (dark purple) and KO (light purple) larvae over time; WT and KO fish have similar median probabilities of posture loss. (G) When summing the minute-by-minute data from panel C, the median time that KO larvae spent displaying normal beat-and-glide behavior was statistically greater than that of WT larvae (p = 0.02). (H): When summing the minute-by-minute data from panel B, the median time that WT larvae spend stationary was statistically greater than the median time KO larvae spend stationary (p = 0.02). (I): When summing the minute-by-minute data from panels D-F, the median time that KO larvae spend displaying seizure-like behavior was statistically greater than that of WT larvae (p = 0.04). (n = 76 WT, n = 36 KO). Statistical significance was determined using the Mann-Whitney test for all panels, with the false discovery rate (FDR) correction applied to Panels B-F, *p<.05, **p<.01, ***p<.001, ****p<.0001.

We next asked whether *camsap3* LOF alters gross larval morphology at baseline. Using the Ramona Kestrel system, we quantified eight anatomical landmarks spanning the head-to-tail axis in wild-type (WT) and knockout (KO) larvae at the same developmental stage (Supplemental Figure 2A). From these landmarks, we derived four primary morphometric outcomes: overall fish length (snout-to-caudal fin), tail length (center-to-caudal fin), head length (snout-to-center), and inter-eye distance (left eye-to-right eye) (Supplemental Figure 2B-E). Across all four measures, we observed no significant differences between genotypes. Total fish length was comparable between WT and KO larvae (Supplemental Figure 2B), as were tail length (Supplemental Figure 2C), head length (Supplemental Figure 2D), and inter-eye distance (Supplemental Figure 2E). Together, these findings indicate that *camsap3* KO larvae do not exhibit overt morphologic abnormalities at this stage, supporting the use of this model for downstream phenotypic analyses without a major baseline body size or morphology confound.

### *Camsap3* mutant larvae show hyperexcitable swimming behavior

We next conducted a proconvulsant provoked swim tracking assay to assesses for seizure like behavior. Behavioral categories came predefined by the Romana Kestrel and are divided into five classifications including stationary, normal beat-and-glide, whirlpool, convulsion, and posture loss. We considered whirlpool, convulsion, and posture loss to be defined seizure-like behavioral states (Figure 4A). Efficacy of the proconvulsant paradigm was confirmed by comparing the behavioral effects of WT PTZ treated control larvae with untreated control larvae. Analyses showed that stationary and normal beat-and-glide behaviors were comparatively similar throughout the recording period (Supplemental Figures 3A-B), while differences in seizure-state probabilities occurred at specific points during the recording window (Supplemental Figures 3C-E). The highest number of convulsions and whirlpool behavior occurred near the beginning of the recording, while posture-loss behavior was observed largely towards the end. (Supplemental Figures 3C-E). Together, these findings suggest that PTZ initially induces clonus-like larval behavior followed by post-ictal like posture loss behavior.

*Camsap3* mutant zebrafish show hyperexcitability, with a pattern similar to that observed in proconvulsant treated larvae compared to untreated larvae. Time resolved analysis revealed differences in the median probabilities of stationary behavior, beat-and-glide, whirlpool, convulsion, and posture loss behavior between WT and KO larvae (Figures 4B-F). The median time KO fish spent displaying normal beat-and-glide behavior was statistically greater than the median time that WT fish spent displaying this behavior (Figure 4G). Similarly, the median time WT fish spent stationary was statistically greater than the median time that KO fish were stationary (Figure 4H). Finally, when summing the minute-by-minute data from panels D-F, the median time that KO fish spent displaying seizure-like behavior was statistically greater than the median time WT fish spent displaying seizure-like behavior (Figure 4I). Together, these findings indicate that, under proconvulsant conditions, *camsap3* KO larvae shift away from stationary behavior and toward seizure-associated swimming states, consistent with increased behavioral hyperexcitability.

### *Camsap3* mutant larvae show increased burst and spike clustering epileptiform

We next tested whether *camsap3* loss of function is associated with abnormal neuronal excitability in vivo using local field potential (LFP) recordings from the optic tectum of 5-7 dpf zebrafish larvae (Figure 5A). Representative tracings showed relatively stable background spiking in WT larvae, whereas KO larvae displayed more frequent high-amplitude burst events and visible spike clustering (Figure 5B).

**Figure 5.**
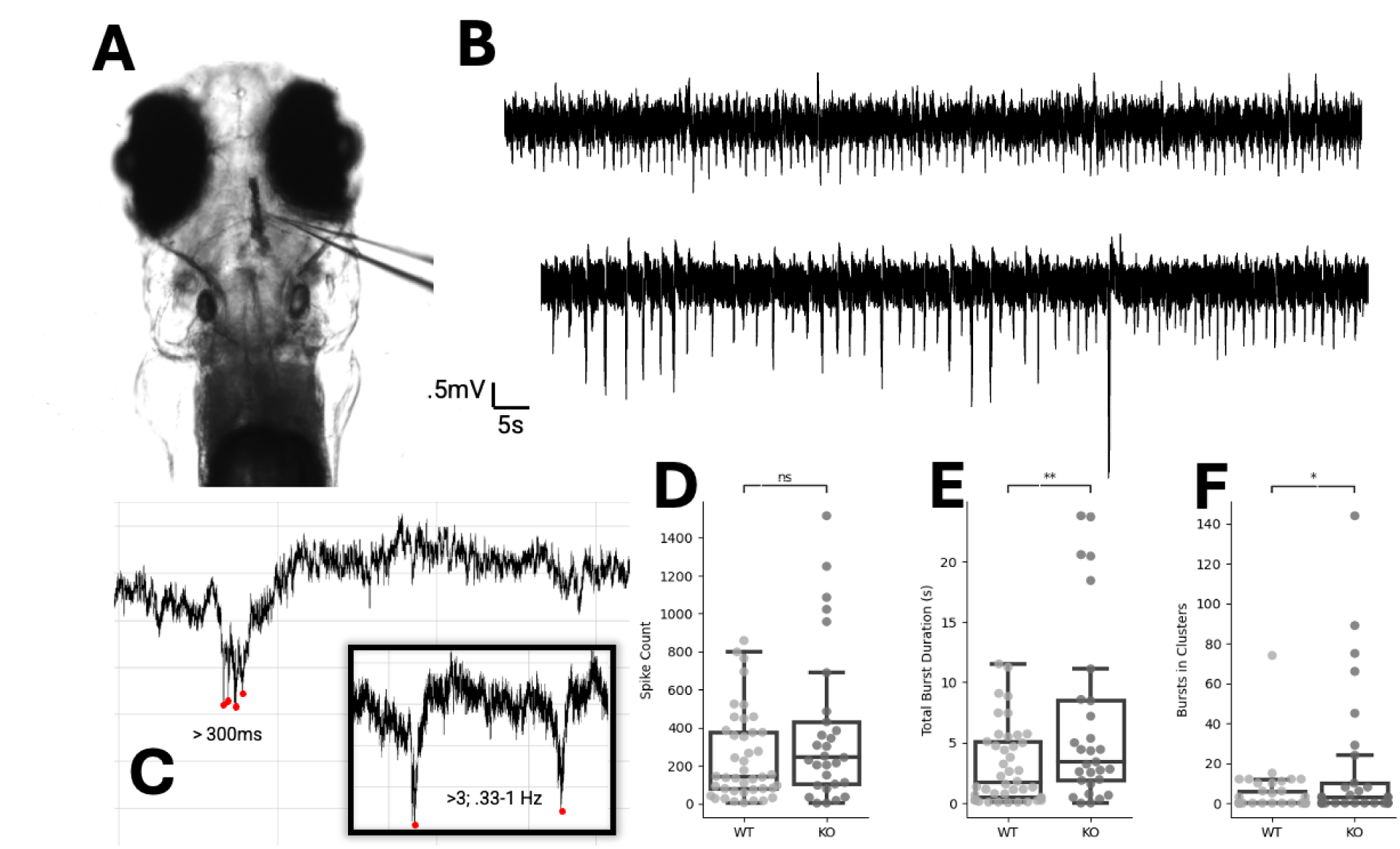
Local field potential recordings in zebrafish larval tectum reveal hyperexcitable network activity. (A) Schematic of single glass recording electrode in the optic tectum of 5- to 7-dpf zebrafish larvae. (B) Representative tracing of normal spiking in a WT larva (top) and abnormal bursting and spike clustering in a *camsap3* KO larva (bottom). (C) Spike bursts were defined as multiple high amplitude discharges greater than three times baseline amplitude in 300 ms. A spike cluster was defined as greater than three spikes in .33-1 Hz. (D) No statistically significant difference in median spike counts is observed between WT and *camsap3* KO fish (p=0.07). (E) Median total burst duration is significantly longer in the *camsap3* KO fish when compared with WT fish (p=0.009). (F) *camsap3* KO fish display a statistically higher median number of bursts in clusters than WT fish (p=0.02) (n = 42 WT, n = 29 KO). Time-resolved statistical significance was determined by Mann-Whitney comparison, with false discovery rate (FDR) correction; ns=not significant, *p<.05, **p<.01, ***p<.001, ****p<.0001.

To standardize event detection, burst events were defined as discharges exceeding three times baseline amplitude within 300 ms, and spike clusters were defined as greater than three spikes within 0.33-1 Hz windows (Figure 5C). ^27^ Using these criteria, we compared WT and KO groups. Analyses showed no significant difference in median total spike count between WT and KO larvae (Figure 5D). In contrast, KO larvae had significantly greater burst-related severity metrics; KO larvae showed longer median total burst duration than WT larvae (Figure 5E) and a higher median number of bursts occurring in clusters (Figure 5F). Together, these findings indicate that while overall spike count was not significantly different, *camsap3* KO larvae exhibit prolonged bursts and increased spike clustering, consistent with a hyperexcitable network phenotype.

### *Camsap3* Loss of Function Reduces Inhibitory Interneuron Populations and Alters Their Spatial Distribution

Given the hyperexcitability observed in mutant *camsap3* larvae, we examined whether *camsap3* loss of function affects inhibitory interneuron abundance and spatial organization in the larval zebrafish brain (Figures 6A-B). To quantify spatial distribution, we reconstructed a 3D tectal surface using Imaris imaging and measured distances from individual interneurons to the tectal midline (Figure 6C).

**Figure 6.**
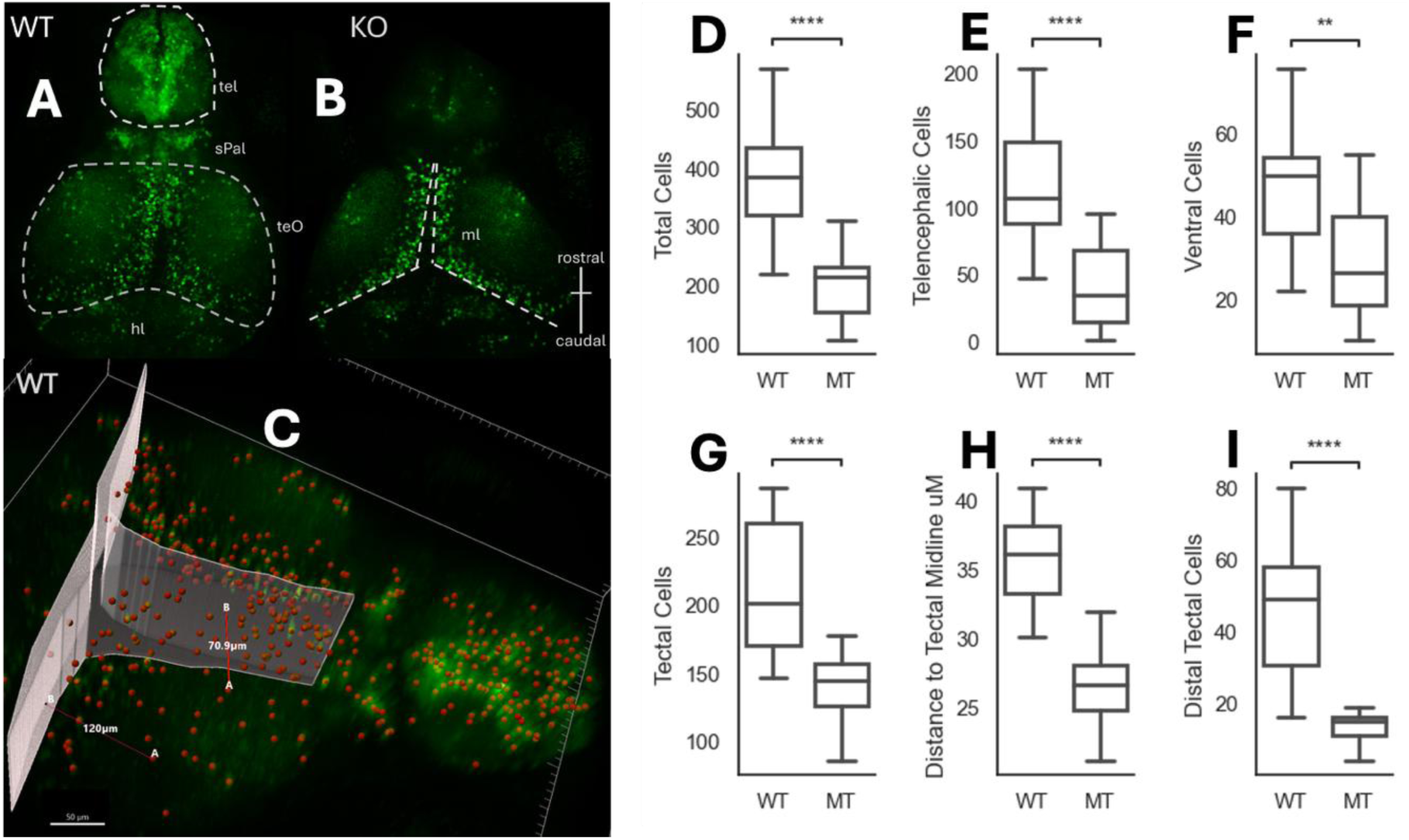
*camsap3* KO leads to mislocalization and decreased total number of inhibitory interneurons. (MT = Mutant/KO). (A) Representative confocal microscopy image visualizing dlx5/6-positive inhibitory interneurons (GFP, green) in the brain of a 7 dpf WT zebrafish. The telencephalon (tel) and the optic tectum region (teO) are marked and outlined in white dotted lines. (B) Representative confocal microscopy image visualizing dlx5/6-positive inhibitory interneurons (GFP, green) in the brain of a 7 dpf KO/Mutant (MT) zebrafish. The midline (mdl) of the optic tectum is highlighted by white dotted lines. (C) Schematic representation of a 3D surface created using Imaris imaging software to quantify the distance of inhibitory interneurons from the midline. Distances between points, or between points and the midline, were measured using lines, such as those shown between points B and A (red lines). (D) Quantification of the total number of dlx5/6-positive inhibitory interneurons found in the tectum, telencephalon, and ventral region of each larva (total cells); *camsap3* KO/MT larvae show a decrease in the mean number of inhibitory interneurons as compared to WT larvae (p<1x10^-4^). (E-G, I) The findings displayed in Panel D (all cells), with a reduction of cells in the mutant vs. WT, reflect comparable differences in mean numbers of inhibitory interneurons in each of the following regions: a separate telencephalic region cell population (E) (p<1x10^-4^), a ventral cell population (F) (p<1.5x10^-3^), a tectal region cell population (G) (p<1x10^-4^), a distal tectal region cell population (I) (p<1x10^-4^). (H) MT/KO fish display a significantly shorter mean distance between dlx5/6-positive inhibitory interneurons and the tectal midline than WT fish, indicating interneuron clustering around the midline in the MT fish (n = 15 WT, n = 16 MT). Statistical significance was determined by unpaired two-tailed Student’s t-tests for all comparisons, *p<.05, **p<.01, ***p<.001, ****p<.0001.

Quantification showed a marked reduction in mean interneuron counts in *camsap3* KO compared to WT larvae (Figures 6D-I). Total dlx5/6+ cell number was significantly decreased in KO fish (Figure 6D), with similar decreases across regional subpopulations, including telencephalic cells (Figure 6E), ventral cells (Figure 6F), tectal cells (Figure 6G), and distal tectal cells (Figure 6I). In addition to reduced cell number, mutant larvae showed altered interneuron positioning: the mean distance of dlx5/6+ cells to the tectal midline was significantly shorter in KO fish (Figure 6H), indicating increased midline clustering. Together, these findings indicate that *camsap3* loss of function is associated with both a quantitative loss of inhibitory interneurons and abnormal inhibitory cell patterning, consistent with disrupted inhibitory circuit development.

### *Camsap3* mutant larvae have increased acetylated α-tubulin in intertectal commissures

We next tested whether *camsap3* loss of function alters acetylated α-tubulin microtubule signal within axon bundles of the intertectal commissure (ITC) (Figures 7A-B). To quantify this effect, midline acetylated-tubulin tracts of the ITC were segmented and reconstructed in 3D using Imaris (Figures 7C-D). Quantitative analysis showed that KO larvae had significantly greater acetylated tubulin signal in both median area (Figure 7E) and median volume (Figure 7F).

**Figure 7.**
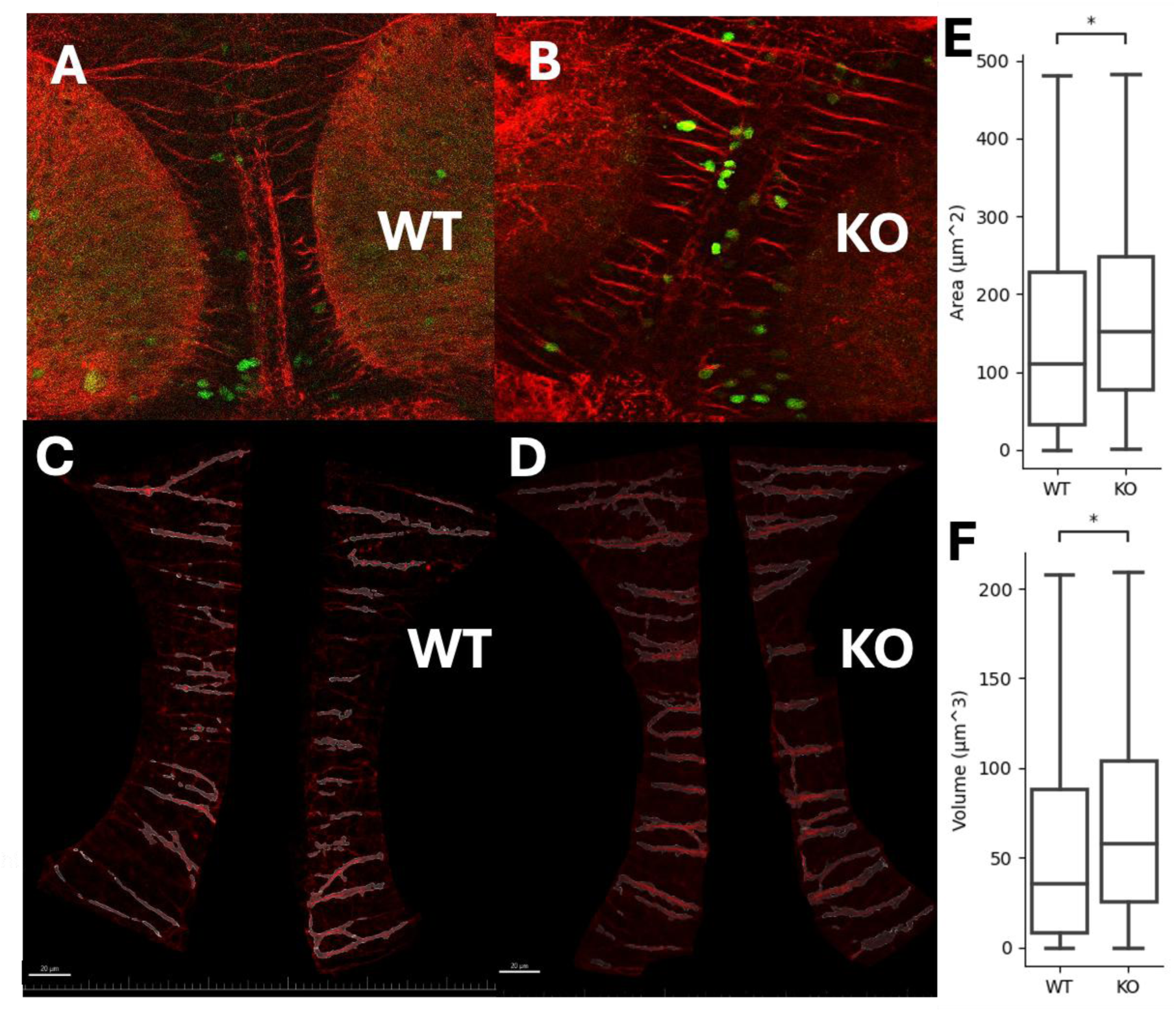
Acetylated alpha-tubulin is more abundant in axon bundles of *camsap3* KO larval intertectal commissure. (A, B) Representative confocal microscopy images of WT and KO larval brains stained with acetylated tubulin (red). (C, D) Schematic maps of tracks of acetylated tubulin along the midline of the intertectal commissure (ITC) in 3D reconstructions of larval brains. (E) *Camsap3* KO fish express acetylated tubulin over a significantly greater area than WT fish (p = 0.04). (F) *camsap3* KO fish express acetylated tubulin within a significantly greater brain volume than WT fish (p = 0.03). (n = 118 WT, n = 111 KO). Significance was determined using the Mann-Whitney test to compare medians, *p<.05, **p<.01, ***p<.001, ****p<.0001.

Together, these findings indicate that *camsap3* loss of function is associated with increased acetylated α-tubulin in the ITC, consistent with altered microtubule organization in axonal commissures due to reduced tubulin deacetylation by *camsap3* in the KO brain.

## DISCUSSION

### Clinical and In Silico Evidence Supports *CAMSAP3* as a Candidate Epilepsy Gene

We evaluated two patients identified by Koh, Smith et al.^9^ and found that both presented with generalized seizures and EEG abnormalities (Figure 1). While Patient 1 showed generalized tonic-clonic seizures, absence seizures, and facial myoclonus with irregular generalized spike-and-slow-wave discharges, and Patient 2 showed generalized spike-wave complexes during absence seizures, their shared generalized EEG signature suggests convergence on a shared network-level phenotype (Figure 1). This clinical convergence is important because generalized epilepsies are typically linked to distributed circuit dysfunction, consistent with the idea that *CAMSAP3* disruption could affect broad neuronal network stability.^13^

Further, we obtained mixed *in silico* predictions (e.g., damaging PolyPhen/CADD results with a less severe SIFT result for one patient). This discordance is not unexpected in rare-variant interpretation and is consistent with guidance from both the American College of Medical Genetics and Genomics and the Association for Molecular Pathology, which treat computational predictions as supporting, rather than definitive, evidence.^14^ In this context, these metrics support the hypothesis that disruptive *CAMSAP3* variants may be functionally consequential but also require further evidence to reach a definitive conclusion.^15^ Importantly, with only two cases, this analysis cannot define penetrance, establish full genotype-phenotype associations, or assign variant type-specific pathogenic certainty on its own. However, the combination of convergent generalized electrophysiology, localization within a biologically relevant *CAMSAP3* domain responsible for neuronal network formation and structure, and partially supportive in silico evidence provides a strong rationale for further investigation. Specifically, modeling *CAMSAP3* variants in vivo in zebrafish allows for the evaluation of epilepsy-related phenomena at the gene level, while experiments in human-derived cells can assess variant-specific functional effects.

### Variant-Specific Modeling in HEK293 Cells Supports a Role for *CAMSAP3* in Microtubule Dynamics

We next modeled the two patient variants ^9^ in an HEK293T overexpression paradigm to determine whether the *in-silico* LOF predictions could be experimentally validated. The lack of GFP signal in Patient 1 (p.K1170RfsX17) is consistent with the premature truncation, while the Patient 2 variant (p.K1023T) results in full-length CAMSAP3 protein, albeit with the single lysine-to-threonine substitution (Figure 2). We next sought to investigate whether variants result in CAMSAP3 LOF by quantifying α-tubulin acetylation. We reasoned that these variants would likely disrupt CAMSAP3 microtubule regulation given their relative positions in the microtubule-binding and CKK domains. The CKK domain of CAMSAP3 has a high affinity for the minus ends, or nucleation sites, of non-centrosomal microtubules. Once bound, CAMSAP3 increases seed stabilization which primes the lattice for acetyltransferase activity. However, LOF experiments in N2a cells and cultured mouse neurons show that siRNA KD of *Camsap3* leads to an increase in microtubule acetylation,^16^ suggesting minus-end-bound CAMSAP3 inhibits αTAT1 acetyltransferase activity. Given this biology, our patient results may seem paradoxical, but the observed effect is due to our paradigm’s supraphysiological levels of protein and the fact that CAMSAP3-mediated acetylation is largely dose-dependent. This is exemplified by *CAMSAP3* overexpression experiments in Caco-2 cells conducted by Pongrakhananon et al ^16^.

These researchers showed overexpression of *CAMSAP3* leads to excessive microtubule binding, not only at the minus end, but also show binding that “streaks” along the microtubule lattice.

Lattice-bound CAMSAP3 stabilizes microtubules, recruiting αTAT1, but inefficiently blocks acetyltransferase activity, resulting in highly acetylated, hyper-stabilized microtubules ^16–19^. Our HEK overexpression experiments corroborate *CAMSAP3*’s dosage sensitivity and show that both LOF and OE paradigms lead to an increase in tubulin acetylation. We showed WT overexpression leads to a large increase in acetylated α-tubulin relative to vehicle controls, while the overexpression of the mutant *CAMSAP3* constructs reduces this effect. Together, these data suggest patient mutations that lead to CAMSAP3 CKK LOF are likely to diminish its function in mediating microtubule acetylation.

### Generation and Validation of a *camsap3* Loss of Function Zebrafish Model

Mouse models of *Camsap3* variants have shown microcephaly due to postnatal ventricular abnormalities and renal tubule cysts leading to enlarged kidneys.^16,20,21^ We did not observe any significant evidence of gross morphological differences between the WT and KO fish assessed (Supplemental Figures 2B-E). Importantly, these data were collected at 6 dpf, the established developmental window for larval zebrafish behavioral assays and approximately equivalent to 6-12 months of age in humans. It remains possible that neurodevelopmental abnormalities emerge later in maturation; for instance, in mouse models of *Camsap3* deficiency, microcephaly was not detected until postnatal day 28, a stage comparable to approximately five years of human development ^16^. Further, enlarged kidneys appeared only in two-year-old mice and appeared normal at postnatal day 21, suggesting that these abnormalities would also likely not be visible when analyzing morphology on 6 dpf zebrafish. ^20^

### Loss of Function Lowers Seizure Threshold and Increases Network Hyperexcitability

We used a validated PTZ paradigm to test seizure-relevant behavior in larval zebrafish (Figure 4). PTZ is a GABA-A receptor antagonist commonly used to induce seizure-like activity in zebrafish by suppressing inhibitory signaling. This treatment results in a stereotypical behavior progression from abnormal hyperactive swimming to convulsive behavior, and ultimately to loss of posture/immobility.^22,23^ In our WT validation comparison (PTZ-exposed versus unexposed), PTZ increased whirlpool behavior across the recording window and increased early convulsive events, while greater posture-loss behavior occurred later in the recording (Figure 4K-M). This temporal sequence supports the interpretation of whirlpool, convulsion, and posture-loss states as seizure-relevant behavioral states. This sequence also strengthens the translational relevance of the assay, as the early hyper-motor and convulsive phases model ictal escalation, whereas later posture loss and reduced movement are analogous to post-ictal suppression after generalized tonic-clonic seizures in humans. Using this established model, we exposed both WT and *camsap3* KO fish to PTZ. When summing the minute-by-minute recording data, *camsap3* KO larvae showed more seizure-associated behavioral states (Figure 4). This result suggests that the seizure threshold in the *camsap3* KO zebrafish is reduced, indicative of a tendency for seizures analogous to human epilepsy. Interestingly, we observed increased total normal swimming behavior and decreased stationary behavior in the KO fish as well (Figures 4G-H). These results suggest that KO larvae spend more time in abnormal motor states overall and exhibit a higher propensity for episodic transitions into overt seizure-like states. Such transitions between normal swimming and seizure-associated states have been described in the literature.^24^ Finally, we detected genotype-driven behavioral differences only after PTZ exposure, not during spontaneous baseline observation. This interpretation is consistent with prior zebrafish literature describing low-dose PTZ as a subthreshold perturbation that can unmask latent epileptogenic susceptibility associated with genetic risk factors, even when baseline phenotypes are subtle or absent.^25^ Additionally, the PTZ proconvulsant model is often necessary because seizure events can be rare relative to the short experimental window. In humans, LOF *CAMSAP3* variants may, in fact, significantly reduce seizure threshold and increase susceptibility to seizures that occur only episodically.

Beyond behavior, epilepsy in humans is diagnosed based on patient history, sometimes supported by direct observation of clinical seizures and EEG evidence of aberrant brain activity indicating a tendency toward seizures or capturing seizures themselves. In larval zebrafish, a local field potential (LFP) recording from the optic tectum, which captures neuronal population-level hyperexcitability, provides an analogous readout to human EEG. We therefore recorded tectal LFPs from larvae and observed that *camsap3* mutant larvae displayed more frequent bursts of LFP spikes in clusters and increased burst duration compared with WT larvae (Figures 5E-F). These features have been associated with epilepsy in other zebrafish models, including models of SCN1A and PCDH19.^26,27^ Therefore, our results demonstrate that *camsap3* gene expression is necessary to establish proper electrical activity balance in the brain and that, with the loss of *camsap3* gene function, the neuronal network is prone to dysregulation in which electrical activity is increased and seizure threshold is lowered. Interestingly, spike counts did not statistically differ between the WT and KO fish, but we posit that this was largely based on how we defined spiking (Figure 5A).

### *Camsap3* disruption alters inhibitory interneuron development and microtubule organization

To examine the impact of *camsap3* loss of function on interneuron abundance and spatial organization, we quantified dlx5a/6a-expressing inhibitory interneurons in the zebrafish tectum and telencephalon (Figure 6). First, we found a significant decrease in the number of dlx5a/6a-expressing inhibitory interneurons in *camsap3* KO larvae compared to WT larvae, suggesting that interneuron reduction could be at least partially responsible for the observed hyperexcitability (Figure 6D). The dlx5/6a reporter is widely used in zebrafish epilepsy studies because dlx-family genes are active early in development and play important roles in interneuron migration and differentiation, and dlx1a, dlx2a, dlx5a, and dlx6a overlap strongly in GABAergic neurons.^28^

Furthermore, we observed alterations, not only in interneuron abundance, but also in the spatial organization of these cells. In the optic tectum, inhibitory interneurons are clustered closer to the midline in the *camsap3* KOs than in the WT fish, suggesting alack of migration of developing neurons away from the midline in the KOs (Figures 6H and I). This result aligns with the core function of the CAMSAP family of genes: to control microtubule dynamics. Microtubules form the cytoskeleton of the nervous system and are crucial for early neuronal development, including migration of the soma. Consequently, defects in microtubule organization or instability directly impair neuronal migration. Similar defects in migration were observed when camsap1 was knocked out in mice ^12^. The literature supports the idea that if microtubules do not properly establish a neuronal network, neurological and neurodegenerative disorders, including epilepsy, can result. ^11^

Previously published findings have shown that CKK-truncated CAMSAP3 expression in cultured mouse neurons leads to hyper-stabilized, highly acetylated microtubules. ^16^ This leads to dysregulated axonal growth as well as axon supernumerary. We aimed to assess whether this dysregulation of tubulin acetylation is conserved in zebrafish and to correlate our CKK LOF patient mutations with our KO phenotype. Consistent with previous results, acetylated alpha-tubulin in KO intertectal commissures had an increased volume compared with WT larvae (Figure 7E and F).

Taken together, our findings suggest that LOF variants in GGE patients lead to loss of *CAMSAP3*-mediated microtubule acetylation. Our zebrafish KO model shows this same acetylation dysregulation, which has been associated with abnormal axon development in mammalian LOF models. Additionally, KO fish have decreased interneuron numbers as well as possible migration defects consistent with tubulinopathies. This decrease and mislocalization of inhibitory interneurons likely lead to hyperexcitability, shown both behaviorally and via LFP, that closely phenocopies seizure manifestations in patients with GGE.

### Limitations

A major limitation of this work is that we have not directly established the relationship between the hyperexcitability phenotype found in our zebrafish mutants and the altered tubulin acetylation dynamics found with *CAMSAP3* loss of function. We show *camsap3* mutant larvae have neuronal migration defects in inhibitory interneurons consistent with tubulinopathies, but we do not show an enrichment of hyper-stabilization supernumerary axons in those neurons.

Such evidence could link the molecular consequences of *camsap3* loss of function with the disruption of network inhibition and the subsequent seizure phenotype. Unfortunately, larval wholemount tubulin staining does not percolate efficiently enough to resolve individual neuronal supernumerary or acetylation defects within the deeper regions of the tectum where dlx5a/6a neurons are found. This is why we were only able to show tubulin acetylation defects within entire axon bundles of relatively superficial commissures of the tectum. Developing a constitutive tubulin reporter zebrafish transgenic crossed with the dlx5/6 transgenic line would allow for measurements of microtubules within inhibitory axons. As of now, the generation of such a transgenic line is beyond the scope of this work but is necessary for understanding the genetic etiology of *CAMSAP3* in GGE.

### Author contributions

Conceptualization: C.M.L., H.Y.K, P.C., and A.P. Methodology: C.M.L., H.Y.K, S.T., L.S., C.M., P.C., and A.P Investigation: C.M.L., Z.S, H.Y.K and H.H. Formal analysis: C.M.L., J.P., and P.C. Resources: A.P Writing – original draft: C.M.L., Z.S., A.P. Writing – review and editing: all authors. Visualization: C.M.L., and P.C. Supervision: C.M.L., P.C., and A.P. Funding acquisition: A.P. Project Administration: C.M.L., P.C., and A.P

### Declaration of Competing Interest

Authors declare no competing interests in relation to the work.

## Acknowledgments

We are grateful and thank the staff of the BCH Zebrafish Aquatic Facility, the Rotenberg Lab and the Experimental Neurophysiology Core (ENC) funded by NIH P50 HD105351, the BCH Cellular Imaging Core; RRID:SCR_026485, and the Ramona Kestrel team for their technical support.

## Funding

AP and the Poduri Lab received support from the National Institute of Neurological Disorders and Stroke (October 2024-July 2025). ST received support from the Rosamund Stone Zander Hansjoerg Wyss Translational Neuroscience Center.

## METHODS

### EXPERIMENTAL MODEL AND STUDY PARTICIPANT DETAILS

#### Zebrafish Husbandry

Adult zebrafish were maintained in 2.8L tanks for no more than 2 years at a density of <30 animals according to standard Animal Resources at Children’s Hospital (ARCH) procedures. 12 hours prior to injection, WT Casper fish were separated by sex (2:1, male: female ratio) into 2L breeding cages. Subsequent embryos were kept at a density of <50 in 100mL of embryo medium (1% methylene blue: fish water) per 100cm petri dish for the first 24 hours of life. From 2-7 dpf fish were grown in fish water only, cleaned daily, and kept in a 27C incubator with a light-dark cycle. All larval experiments were conducted on or before 7 dpf prior to yolk absorption to avoid feeding confounds later in development. If larvae were intended to be grown to adulthood, they were placed in 1.8L tanks prior to 6 dpf and reared by the ARCH aquaculture team.

#### Cells

Human embryonic kidney cells (HEK293T) were grown in culture medium (0.22 µm vacuum filtered Dulbecco’s modified Eagle medium containing 10% fetal bovine serum) and aliquoted in culture medium containing 1% DMSO and stored at -125C. Cells were maintained in an incubator at 37C and passaged no more than twice before experimentation.

#### Human Participants

Boston Children’s Hospital (BCH) patient variants were identified in a previously published cohort study of individuals with unexplained epilepsies. Patients or guardians were recruited for the study between August 2018 and May 2023, and the study was approved by the BCH Institutional Review Board. Variants were annotated to RefSeq mRNA transcript NM_001080429.2 and NCBI RefSeq protein accession NP_001073898.1. Previously reported variants included Patient 1: GRCh37(hg19) chr19:7682528AG>A; c.3507del; p.(Lys1170ArgfsTer17) and Patient 2: GRCh37(hg19) - chr19:7680480A>C; c.3068A>C; p.(Lys1023Thr).

### METHOD DETAILS

#### Zebrafish Model Generation

C*amsap3* CRISPR guide (Supplemental Figure 1) was chosen based on the highest specificity and efficiency predicted by the CHOPCHOP design tool (https://chopchop.cbu.uib.no/) and then generated by Synthego. Guides were diluted and aliquoted in RNase/DNase free ddH₂O to 1000 ng/nl and stored at -80C. 2nl of CRISPR complexes (500ng/nl guides : 500ng/nl of Cas9 protein) were injected via a forced-air microinjector at the one-to two-cell stage. Injected larvae were grown to sexual maturity (>3 months) and were genotyped by PCR/Sanger sequencing (Genewiz) using fin clips. Mutant F0 fish containing the truncating mutation were outcrossed to Casper or tgn(gfp:dlx5/6; rfp:vglut) WTs. F1 fish were then generated and incrossed for stable F2 experiments. All subsequent generations alternated between WT outcrosses and genotyped incrosses.

#### Behavior

Larvae from het x het crosses were grown to 6 dpf and pipetted in 100 µL of fish water per well of a 96-well screening plate. Plates were placed in a Ramona Optics Kestrel and allowed to acclimate for 10 minutes. The plate was run in the high resolution “behavior mode” at 160 fps for 30 minutes for the spontaneous trial. 100 µL of 10 mM PTZ fish water was then added to each well for a final concentration of 5 mM. The provoked trial was then run for an additional 30 minutes. Data were then exported, and ontologies were processed and categorized using the built-in MCAM classifications recently developed ^22^and reported^29^.All post-processing was done with GraphPad and Python.

#### Electrophysiology

Larvae were grown to 5-7 dpf from het x het crosses and paralyzed in 2 mg/ml α-bungarotoxin for 10 minutes or until loss of touch response. Two larvae per recording session were mounted on glass stages in 2% low-melting-point agarose heated to 37C. Glass microelectrodes backloaded with M were generated using a heated microelectrode puller to a resistance of 1-7 MΩ. Electrodes were guided to a position slightly offset from the midline within the larval optic tectum via a micromanipulator. Voltage clamp recordings were conducted using pClamp 12 software and filtered at lowpass:1kHz (3 dB; eight-pole Bessel) and high-pass: 0.1-0.2 Hz.

Voltage was also amplified with an Axopatch 200B and digitized at 5-10 kHz using a Digidata 1440A. Fish were continually monitored for heartbeat and blood flow throughout the 30 minute spontaneous recording and were discarded if they died. Data were analyzed using a modified version of previously described custom Python code: (https://doi.org/10.5281/zenodo.11165681).

#### Transfection and Immunocytochemistry

Human CAMSAP3 vector was mutagenized using standard Agilent procedures. Primers (Supplemental Table 1) were designed for each patient mutation using the Agilent QuickChange primer design web app (https://www.agilent.com/store/primerDesignProgram.jsp;?srsltid=AfmBOoq0ukVP1JHopF-acAMBOTfhRCoePVa42BWUOahYgY6Dq0Hoo1Cy). Constructs were amplified via transformation with chemically competent TOP10 cells and isolated via Qiagen miniprep. HEK293T cells were grown in poly-L-lysine-coated 6-well plates to 70% confluence and transfected (5 µL Lipofectamine 2000: 2,500 ng total DNA per reaction) in serum-free medium under standard kit procedures. The transfection reaction was incubated for 36 hours before transferring cells to 8-well chamber slides for 12 hours. Slides were fixed with 4% PFA in PBS overnight at 4C and then washed (3x for 5 minutes) with TBS-Tween 20 at room temperature. Cells were then incubated in blocking buffer (1% DMSO, 2% NDS, and 1% BSA in PBT) for 1 hour at room temperature, followed by the addition of primary antibody (alpha-tubulin rb 1:1000) and staining overnight at 4C. Stained cells were then washed, incubated in blocking buffer with secondary antibody (Alexa Fluor 405 rb 1:1000) for 1 hour at room temperature, and washed again. Chamber wells were then removed, and slides were mounted with a droplet of mounting medium and a coverslip. Imaging was conducted using a Zeiss 700 confocal microscope and analyzed using Imaris version 11.0.

#### Whole Mount Zebrafish Analysis

Both Casper and tgn(gfp:dlx5/6; rfp:vglut) larvae were sacrificed on ice at 6 dpf and fixed in 4% PFA in PBS overnight at 4C for respective analyses. The PFA wash consisted of 3x washes with TBS-T for 5 minutes at room temperature. Larvae with endogenous fluorescence were mounted intact in 2% low-melting-point agarose and immediately imaged, and brains were extracted using standard procedures. Brain isolates were incubated in 150 mM Tris-HCl, pH 8, for 5 minutes at room temperature and 15 minutes at 70C for antigen retrieval. Isolates were then washed and permeabilized in 0.05% trypsin-EDTA at 37C for 45 minutes. Larval brains were then washed again and incubated in blocking buffer (2% NDS, 1% BSA, and 1% DMSO in PBS-T) for 1 hour at room temperature. Staining consisted of primary incubation (1:1000 alpha-tubulin ms (Sigma) in blocking buffer overnight at 4C), a subsequent wash, and secondary incubation (1hour at room temperature Alexa Fluor 568 ms), followed by another 3x wash. Stained brains were then mounted on slides with mounting medium containing DAPI (Invitrogen) and a thin outline of vacuum grease to keep the weight of the coverslip off the sample and to maintain brain structure. Imaging was conducted using a Zeiss 700 confocal microscope, and analysis was done using Imaris version 11.0. Axon fibers were automatically detected using Imaris machine-learning module, and dlx-positive neurons were automatically detected using the seeding module.

## Notes

### Competing Interest Statement

The authors have declared no competing interest.

