## Supplement for "CAMSAP3 loss of function models suggest causative role in generalized genetic epilepsy"

**Table 1: Primers**

| Sequence | Primer | Purpose |
|---|---|---|
| TAGGTCCATCGTTAGGTTTGCT | z_camsap3_genomic_F | Fish KO detection in DNA |
| ACTAACTGCAAACGGTGTCCTT | z_camsap3_genomic_R | |
| CCCGCTGGTACTGGAAACTT | z_camsap3_cDNA_1_F | Fish KO detection in cDNA |
| AGACAGCGGACGACTAAGTG | z_camsap3_cDNA_1_R | |
| CAGGAGGTGGAGAAGGAACA | h_camsap3_vector_cterm_F | Vector validation for HEK cell experiments (missense and truncating mutations) |
| TTGGGAGTGGTGGGTTTCTT | h_camsap3_vector_cterm_R | |
| GGCCGAGTTGTTCATGTGTT | h_camsap3_vector_nterm_F | Vector validation for HEK cell experiments (WT) |
| GCACTGTGGATGATCTGCAG | h_camsap3_vector_nterm_R | |

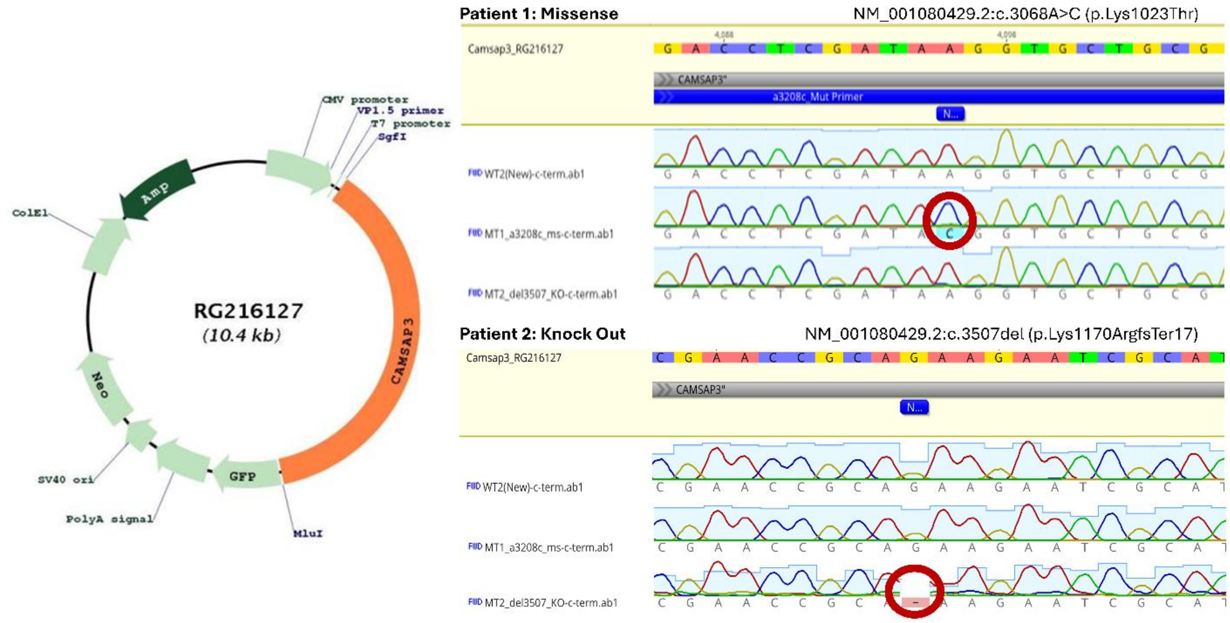


**Figure 1**. **Plasmid map (adapted from OriGene) and patient mutation validation.**

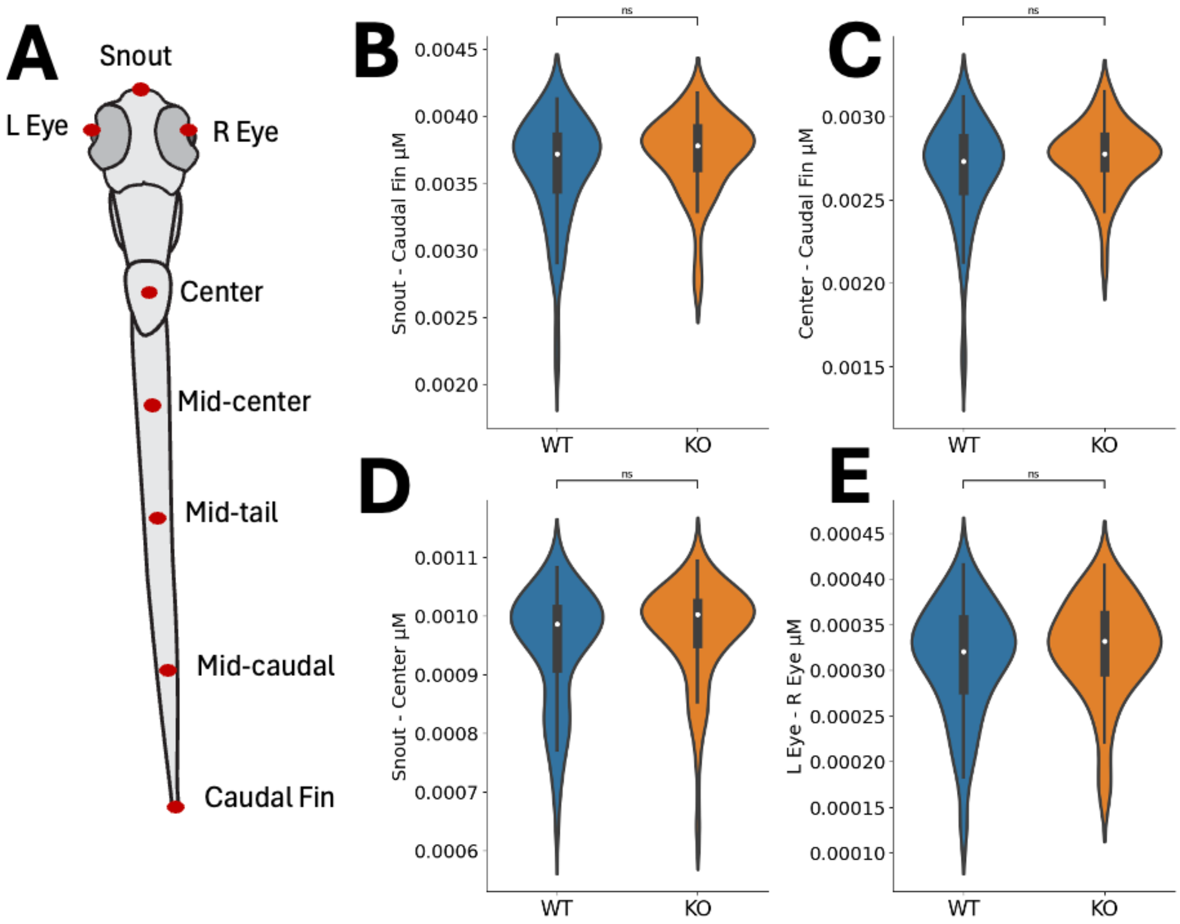


**Figure 2. Camsap3 KO larvae have normal gross morphology.** (A) Using the Ramona Kestrel system, zebrafish larvae were measured at eight points from the eyes and snout to their caudal fin. No morphological differences between WT (blue) and KO fish (orange) were observed, including in median fish length (B) ($p=0.13$), tail length (C) ($p=0.15$), head length (D) ($p=0.12$), and inter-eye distance (E) ($p=0.28$) ($n = 76$ WT, $n = 59$ KO). Statistical significance was determined by Mann-Whitney, ns=not significant.

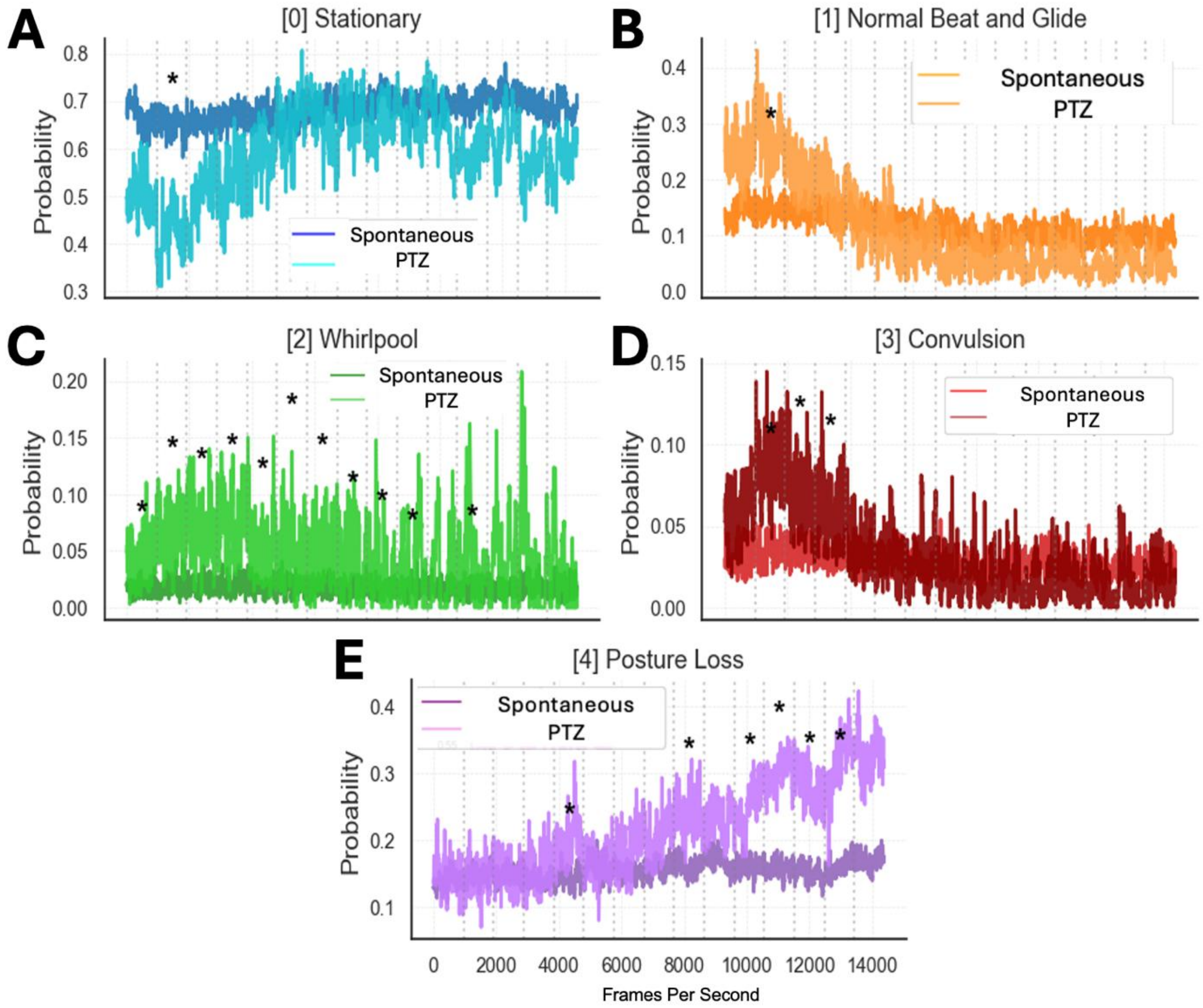


**Figure 3. Impact of the proconvulsant drug PTZ (pentylenetetrazol) on WT larvae swimming patterns compared to untreated (spontaneous) WT larvae over time. (A and B):** Other than for one minute at the start of recording, there were no significant differences in median total stationary and total normal beat-and-glide swim probability between PTZ-treated and untreated fish over time. **(C, D, and E):** Differences in median probability of whirlpool swimming, convulsions, and posture loss over time were observed between PTZ-treated and untreated fish; PTZ-treated fish showed a statistically higher median probability of whirlpool throughout minutes 1 through 9 and 11 (**C**), a statistically higher median probability of convulsion during minutes 1 through 3 (**D**), and a statistically higher median probability of posture loss during minutes 4, 8, and 10 through 13 **(E).** (n = 76 for spontaneous and PTZ). Time-resolved statistical significance was determined by Mann-Whitney with false discovery rate (FDR) correction, $*p<.05$.
